# FGF21 Analogue PF-05231023 on Alcohol Consumption and Neuronal Activity in the Nucleus Accumbens

**DOI:** 10.1101/2024.12.22.629996

**Authors:** Bart J. Cooley, Cassandra V. Occelli Hanbury-Brown, Eun A. Choi, Willow A. Heller, Alyssa W. Lim, Andrew J. Lawrence, Paul S. Haber, Gavan P. McNally, E. Zayra Millan

## Abstract

Fibroblast growth factor 21 (FGF21) is a liver-derived hormone known to suppress alcohol consumption in mice and non-human primates. However, the role of FGF21 in modulating environmental and behavioural factors driving alcohol consumption—such as cue-driven responses and effortful actions to obtain alcohol—and its effects on neural activity related to consumption, remain unclear. Here, we evaluated the impact of PF-05231023, a long-acting FGF21 analogue, across multiple dimensions of alcohol consumption and motivation. PF-05231023 reduced alcohol intake and preference in a dose-and sex-specific manner; diminished approach behaviours following an alcohol but not sucrose cue; and decreased lever-pressing under a progressive-ratio schedule, both alone and when combined with the GLP-1 agonist Exendin-4. Additionally, PF-05231023 altered the microstructure of alcohol consumption by shortening drinking bouts and increased the recruitment of nucleus accumbens (Acb) neurons associated with bout termination. These findings demonstrate that PF-05231023 broadly suppresses alcohol-motivated behaviours and that targeting FGF21 signaling in combination with GLP-1 agonists may enhance therapeutic efficacy. Mechanistically, the observed reductions in alcohol consumption following PF-05231023 appear to involve diminished alcohol palatability and modulation of neuronal activity from distinct subsets of Acb neurons.

## Introduction

Alcohol use and related harms are a leading cause of global morbidity (1). Medications for alcohol use disorder (AUD) like naltrexone, nalmefene, acamprosate, and topiramate are available, but their effectiveness varies widely, leaving a significant gap in treatment options (2, 3). Emerging pharmacological targets include peptides that promote metabolic health (glucose homeostasis, insulin sensitivity) and energy balance, such as those secreted by the gut (e.g., GLP-1, ghrelin, (4–7)), brain (e.g., orexin, (8–10)), and adipose tissue (e.g., leptin, (11)). These peptides exploit shared mechanisms between the regulation of alcohol-motivated behaviour and satiety pathways (11, 12), but there is emerging interest in fibroblast growth factor 21 (FGF21)—a liver-secreted endocrine peptide that regulates macronutrient preference (13–15)—as a promising new target, complementary to the actions of other energy-balancing peptides (16, 17), for reducing alcohol consumption.

FGF21 is induced in the liver in response to various metabolic stressors, including alcohol (18–20). It signals via binding to an FGFR1c/β-klotho (KLB) cell surface receptor complex (21, 22). While it acts at multiple peripheral tissues, it can also act centrally to bias macronutrient choice (15), indicating its potential to directly regulate appetitive behaviour, though the extent of this regulation for conditions involving alcohol consumption are not well characterised. Three lines of evidence support the role for FGF21 in the regulation of alcohol consumption. First, overexpression of FGF21 in transgenic mouse lines significantly reduces alcohol preference (23) while pharmacological administration of FGF21 or its analogue, PF-05231023, reduces alcohol consumption in mice and alcohol-preferring vervet monkeys (24). Second, in large-scale genome wide association studies in humans, single nucleotide polymorphisms (SNPs) in *FGF21* and *KLB* regulatory genes were significantly associated with alcohol consumption and AUD risk (25–27). Finally, disruption of central FGF21 signalling via brain-specific knock out of the obligate FGF21 coreceptor, KLB, increased alcohol consumption in mice (27).

The central actions of FGF21 on alcohol consumption have been localised to an amygdala-ventral striatal pathway involving KLB+ neurons in the basolateral amygdala (BLA) that project to the nucleus accumbens (Acb, (24)), consistent with prior studies implicating this pathway in regulating alcohol intake and cue-driven conditioned responses in the presence of alcohol-paired cues (28). The Acb in particular is centrally positioned to rapidly suppress intake of orally-ingested rewards, with previous studies showing that stimulation of the Acb or its projections inhibits consumption (29, 30). Conversely, pauses in the activity of Acb cells are required to initiate and maintain consumption (29, 30), supporting its role as a gateway for consummatory behaviour. Whether and how FGF21 acts directly on individual Acb cells to diminish these pauses remains unknown.

The evidence implicating FGF21 or its analogues over the regulation of alcohol consumption is important but is broadly limited to consumption *per se*. The influence of FGF21 on precursors to drinking, including approach triggered by alcohol-associated cues and instrumental actions required to obtain alcohol remain unknown. We addressed these here and first show that PF-05231023 reduces voluntary alcohol consumption and preference in male but not female mice in a dose-dependent manner. PF-05231023 also attenuates responses following the presentation of an alcohol-but not sucrose-predictive cue and reduces the motivation to seek alcohol on an instrumental progressive ratio test. To understand how PF-05231023 suppresses alcohol consumption we examined the microstructure of licking in alcohol drinking mice while recording the activity of individual neurons in the Acb. We found that PF-05231023 reduced the palatability of alcohol and modulated specific subsets of Acb cells associated with initiating and terminating consumption.

## Materials and Methods

### Animals

All procedures were approved by the Animal Care and Ethics Committee at UNSW Sydney. All procedures conformed to ARRIVE guidelines, C57BL/6J mice (OzGene, Perth, Australia) were single-housed on a 12:12-hour light/dark cycle (lights on at 07:00 for all experimental studies). During homecage alcohol consumption studies, mice had *ad libitum* access to food and water. For Pavlovian and instrumental studies, mice were maintained at 85% of their baseline body weight under food restriction. During lickometer studies, mice were mildly water-restricted by removing water bottles ∼5 hours prior to testing. Behavioural testing occurred during the light phase.

### Surgeries and viral injections

To monitor dynamic neural activity in Acb, an adeno-associated virus encoding a calcium sensor (AAV9-syn-GCaMP7f) was injected into the Acb, followed by a gradient refractive index lens (GRIN) implanted above the injection site (details in Supplementary Methods).

### Drugs

Alcohol solutions (15%) were prepared by diluting 100% ethanol with tapwater. PF-05231023, a long-acting FGF21 analogue (AdooQ Biosciences, CA, USA), and Exendin-4 (3-39; Bachem), a long-acting GLP-1 analogue, were dissolved in saline (0.5% v/v DMSO in Experiments 1 and 4 only) using three cycles of 15-minute warm water baths (36°C) and sonication. Doses were based on previous studies in alcohol-drinking mice and non-human primates (24). Vehicle was saline (0.5% v/v DMSO in Experiments 1 and 4 only). Drugs were administered intraperitoneally at 10ml/kg. Injections occurred immediately, 60 minutes or 2-4 hours prior to 24hr homecage access, cue-approach/lever pressing, and lickometer tests, respectively.

### Cellular-resolution calcium imaging

Calcium imaging of Acb neurons during alcohol consumption was synchronized with lick-event timestamps in mice (N=8) with correctly placed virus and GRIN lens (Supplementary Figure, S2). Detailed methods are provided in the Supplementary Methods.

### Procedure

Detailed methods are provided in the Supplementary Methods.

#### Experiment 1. Effect of PF-05231023 on homecage alcohol consumption

Mice received 3 weeks of 24-hour intermittent alcohol access. Test occurred between Weeks 4-8. Mice received vehicle, 1mg/kg, 3mg/kg or 10mg/kg PF-05231023 at the start of each week in a within-subjects Latin square design to control for order and time effects. Intake was measured at 24hrs, 72hrs and 120hrs post treatment.

#### Experiment 2. Effect of PF-05231023 on response to alcohol and sucrose cues

Mice were trained to associate auditory cues with alcohol (CS1) or sucrose (CS2). They were trained in two blocks: sucrose cue training (CS+, CS-; 15 trials/cue/session; 16 sessions, daily) then alcohol cue training (CS+, CS-; 10 trials/cue/session; 16 sessions, daily). Between the two blocks they received 3 weeks of overnight intermittent access to alcohol. They were tested under 10mg/kg PF-05231023 and vehicle in the following manner: CS1 treatment 1→48hr washout→CS1 treatment 2→6 day washout→CS2 treatment 1→48hr washout→CS2 treatment 2, where CS1 and CS2 are sucrose and alcohol cues (counterbalanced order); and treatment 1 and treatment 2 are drug and vehicle (counterbalanced order). Test comprised 20 trials (10 CS+, 10 CS-, pseudorandom order, VI range 100-120s). Subsequently, mice were retrained with the sucrose CS+ (3 sessions) and alcohol CS+ (3 sessions). They were then tested in the same manner for the effects of the lower dose, 3mg/kg PF-05231023 (Fig.2A).

#### Experiment 3. Effect of PF-05231023 on progressive ratio

Male mice were tested on a PR schedule under one of four treatments, counterbalanced in a repeated measures Latin-square design: vehicle, PF-05231023 (3mg/kg), Exendin-4 (Ex4; 2.4µg/kg), and combined [PF-05231023 (3mg/kg) + Ex4(2.4µg/kg)]. Mice received one PR test per week. They were retrained on RR10 at 2 and 4 days post-test to maintain a stable level of lever pressing across testing weeks. They were then tested in a similar manner under one of three treatments: vehicle, PF-05231023 (10mg/kg), and combined [PF-05231023 (10mg/kg) + Ex4(2.4µg/kg)].

#### Experiment 4. Effect of PF-05231023 on cellular transients in Acb during alcohol consumption

Male mice had 20 min access to alcohol from a lickometer spout in an experimental chamber. Recordings of Acb task-related cellular activity were acquired on Days 5-8. They received saline to acclimatise to the injection procedure on Day 5, then Vehicle and PF-05231023 (10mg/kg) on Days 6 and 7, respectively.

### Data analyses and statistics

We analysed behavioural and calcium imaging data using custom scripts in R and Python, respectively. Repeated-measures comparisons between treatment groups were analysed using a planned set of orthogonal contrasts with the PSY statistical program (Bird, 2004), unless otherwise specified. Specific analyses are described in Supplementary Methods.

## Results

Across experiments mice consumed stable and moderate levels of alcohol during homecage two bottle choice (Supplementary Table 1).

### Experiment 1: PF-052131023 reduces alcohol consumption

We first assessed the dose-response effect of PF-05231023 on alcohol consumption in male and female mice. We excluded two mice (n=2 males) that exhibited low and unstable alcohol consumption prior to test. There was a dose-and sex-selective effect of PF-05231023 (Fig.1). In male mice there was a U-shaped dose-response curve when consumption was measured 24 hours after injection. Specifically, alcohol intake and preference was significantly reduced following 3mg/kg PF-05231023 (Drug quadratic trend: alcohol intake, F(1,9)=9.381, p=0.014; alcohol preference, F(1,9)=8.955, p=0.015, Figs.1A and C). There was no main effect of the drug compared to vehicle at 24 hours post-injection (Drug main effect: alcohol intake, p=0.228; alcohol preference, p=0.223), and there were no differences in alcohol intake and preference under 1mg/kg and 10mg/kg PF-05231023 (Simple effect: alcohol intake, p=0.197; alcohol preference, p=0.216). PF-05231023 also attenuated alcohol preference but not intake 72 hours after treatment, which did not follow a U-shaped curve (Drug main effect: alcohol preference, F(1,9)=5.749, p=0.040; alcohol intake, p=0.217; Drug quadratic trend: alcohol intake, p=0.428; alcohol preference p=0.491, Fig.1C). There was no effect of PF-05231023 on preference or intake at 120 hours after treatment (all contrasts: intake, p>0.087; preference, p>0.214). Critically, the effect of treatment was not due to a non-specific decrease in liquid consumption because there was an inverse U-shaped dose-response effect of PF-05231023 on water consumption 24 hours following treatment (Drug quadratic trend: F(1,9)=6.584, p=0.030; Drug main effect: p=0.440; 1mg/kg or 10mg/kg simple effects: *p*s>.087; Fig.1B). This effect did not persist at 72 and 120 hours post-treatment (all contrasts, p>0.087).

**Figure 1.**
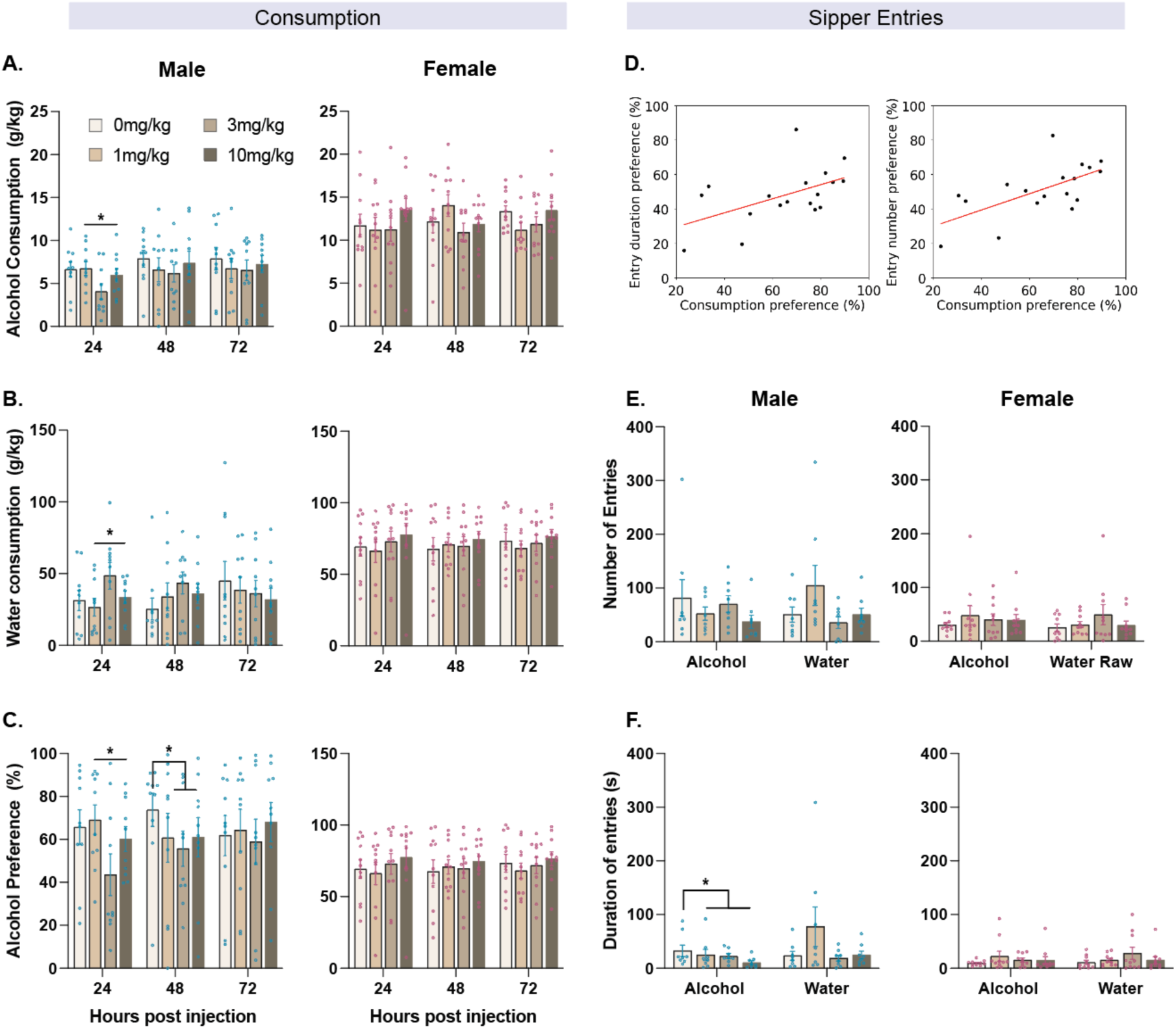
Effects of PF-05231023 (0mg/kg, 1mg/kg, 3mg/kg, 10mg/kg) on alcohol consumption. **(A)** Alcohol consumption, **(B)** Water consumption and **(C)** Percentage alcohol preference in male and female mice at test. **(D)** Scatterplot of consumption and entry duration / number preference on the last three days of intermittent access prior to test. **(E)** Number of entries and **(F)** duration of entries in male and female mice at test. Data are means±SEM. *p<0.05. N = 23 (male n=12).

In females, there was no effect of PF-05231023 on measures of alcohol (all contrasts: alcohol intake, p>0.120, alcohol preference, p>0.559) or water consumption (p>0.281). We also did not find any effect of the drug on body weight of animals (all contrasts: females p>0.300, males p>0.80). Together these findings suggest that PF-05231023 has a dose-dependent effect on alcohol consumption and preference in male but not female mice.

We next analysed sipper entry data to determine whether PF-05231023 altered homecage drinking behaviours more broadly. Sipper entry tracked consumption of alcohol and water (preference scores) because preference on the last three days of intermittent access prior to test correlated with frequency (R^2^=0.40, p=0.01) and duration of entries (R^2^=0.27, p=0.03, intermittent access days 7-9) (Fig.1D). We focused analyses on the first day of test (24hr post injection) given that treatment effects were largest at this timepoint. Our data indicated a dissociation in the effects of PF-05231023 on approach to the sipper and consumption of liquids in male but not female mice. In males under vehicle, sipper entry and duration preference linearly tracked consumption preference (Entry preference, R^2^=0.588, p=0.016; Duration preference, R^2^=0.619, p=0.012) whereas under any tested dose of PF-05231023 there was no significant linear relationship between consumption preference and approach to the sipper (1mg/kg: Entry preference, R^2^=0.385, p=0.056; Duration preference, R^2^=0.217, p=0.175; 3mg/kg: Entry preference, R^2^=0.265 p=0.128; Duration preference, R^2^=0.042, p=0.570;10mg/kg: Entry preference, R^2^=0.058, p=0.502; Duration preference, R^2^=0.036, p=0.598). We did not find this effect in females, although the linear relationship varied across doses of the drug and vehicle (Vehicle: Entry preference, R^2^=0.335, p=0.133; Duration preference, R^2^=0.246, p=0.212; 1mg/kg: Entry preference, R^2^=0.436, p=0.075; Duration preference, R^2^=0.548, p=0.036; 3mg/kg: Entry preference, R^2^=0.862, p=0.002; Duration preference, R^2^=0.862, p=0.003; 10mg/kg: Entry preference, R^2^=0.396, p=0.094; Duration preference, R^2^=0.382, p=0.102). This suggests a dissociation between the effects of PF-05231023 on consumption and sipper approach. Interestingly under drug, male (Main effect: F(1,9)=7.034, p=0.026, quadratic trend, p=0.623) but not female mice (Main effect: p=0.302, quadratic effect: p=0.558) spent more time at the alcohol sipper (Fig.1E). These effects did not extend to other measures because there was no significant effect of the drug on the number of alcohol sipper entries (females: p=0.637, males: p=0.115, Fig.1F) or average entry duration (females: p=0.310, males: p=0.935) and no quadratic trend on either measure (Entries: females: p=0.097, males: p=0.820; Average duration: females: p=0.976, males: p=0.448). We found no effect of PF-05231023 at the water sipper on the number of entries, duration of time, or average entry duration (All contrasts, females: *p*s>0.102; males: *p*s>0.215).

### Experiment 2: PF-05231023 reduces alcohol-cue approach without affecting sucrose-cue approach

Next, we assessed the effects of PF-05231023 on conditioned approach to an alcohol-predictive stimulus. We first trained mice with sucrose-cue approach. As expected, mice showed significant conditioned approach during the sucrose cue (CS+>CS-, frequency, F(1,9)=17.622, p=0.002; probability, F(1,9)=137.143, p<0.001). They then received intermittent homecage access to alcohol followed by training with an alcohol-paired cue (Fig.2A). However, despite equivalent training, mice did not acquire conditioned approach during the alcohol cue (last three days of training: frequency: p=0.222; probability: p=0.717). However, they showed heightened responses to both CS+ and CS-compared to baseline (F(1,10)=7.22, p=0.019) and distinguished cues based on alcohol availability at CS+ offset (Figs.2C-D). For this reason, we applied separate and different analyses to assess the effects of PF-05231023 on responses to sucrose and alcohol cues.

**Figure 2.**
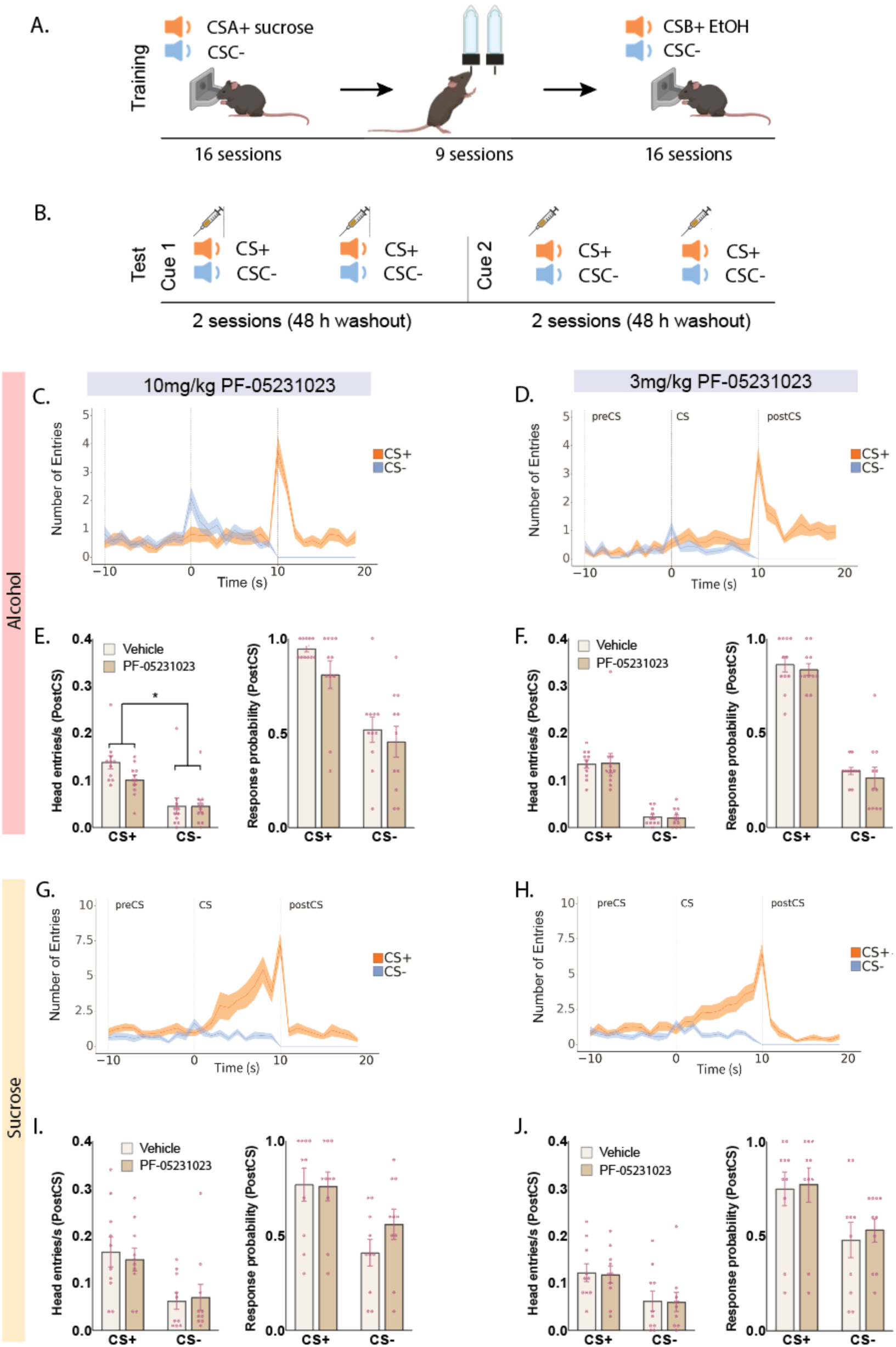
Effects of 3mg/kg and 10mg/kg PF-05231023 on approach to alcohol and sucrose paired cues **(A)** Schematic of training and **(B)** test procedure of conditioned approach to alcohol and sucrose paired cues. Cue 1 and Cue 2 are alcohol and sucrose cues counterbalanced across mice. Treatment order was counterbalanced. **(C)** Magazine entries during alcohol CS+ and CS-trials separated into minute time bins including the preCS (-10-0 seconds), CS (0-10 seconds) and postCS (10-20 seconds) under 10mg/kg and **(D)** 3mg/kg PF-05231023. **(E)** Frequency and probability of magazine entry during presentation of alcohol rewards (postCS) under 10mg/kg PF-50231023. (**F)** Frequency and probability of magazine entry under 3mg/kg PF-05231023 during alcohol postCS+ and postCS-. **(G)** Magazine entries during sucrose CS+ and CS-trials separated into minute time bins including the preCS (-10-0 seconds), CS (0-10 seconds) and postCS (10-20 seconds) under 10mg/kg PF-05231023 and **(H)** 3mg/kg PF-05231023. **(I)** Frequency and probability of magazine entry during presentation of sucrose rewards (postCS) under 10m/kg PF-05231023 and 3m/kg PF-05231023. Data are means±SEM. *treatment x cue interaction, p<0.05. N=10 (male: n=6).

On alcohol cue tests (Figs.2C-2F), approach behaviour was significantly elevated during the CS (Cue main effect: F(1,10)=15.454, p=0.003) confirming the learned significance of these cues. Approach responses during alcohol delivery at CS+ offset was specifically elevated relative to control CS-(frequency: F(1,10)=172.153, p<0.001; probability, F(1,10)=230.440, p<0.001). While there was no effect of PF-05231023 on approach during the CS (10mg/kg: p=0.514; 3mg/kg: p=0.591), PF-05231023 significantly reduced the frequency of approach during alcohol delivery at the highest dose of 10mg/kg (treatment x cue, frequency: F(1,10)=6.817, p=0.026; probability: p=0.333, Fig.2E). There were no effects following the 3mg/kg dose (frequency: treatment x cue, p=0.866; probability: treatment x cue, p=0.905; Fig.2F).

On sucrose cue tests, mice discriminated sucrose CS+ from control CS-cues, showing elevated approach during the CS+ (frequency: F(1,9)=5.814, p=0.039; probability: p=0.096) and during sucrose delivery at CS+ offset (frequency: F(1,9)=5.912, p=0.038; probability: F(1,9)=8.69, p=0.016). Critically, PF-05231023 did not affect conditioned approach during the sucrose-predictive cue (10mg/kg or 3mg/kg; all *p*s>0.072) or sucrose delivery at either dose (10mg/kg or 3mg/kg; all *p*s>0.129; Figs.2G-J).

Together these findings show that PF-05231023 dose-dependently attenuates cue-elicited responses during alcohol availability without affecting similar responses to a sucrose-paired cue. Thus, PF-05231023 may impart selective effects on alcohol-motivated behaviour.

### Experiment 3: PF-05231023 attenuates alcohol seeking under a progressive ratio schedule

Next, we assessed the effects of PF-05231023 on motivation to respond for alcohol using a progressive ratio schedule (Fig.3A). Given recent interest in dual FGF21/GLP-1 agonist approaches (16, 17) we also tested PF-05231023 in combination with the GLP-1 agonist, Exendin-4. We used a low dose of Exendin-4 previously shown to minimise off-target effects on locomotor performance (6).

**Figure 3.**
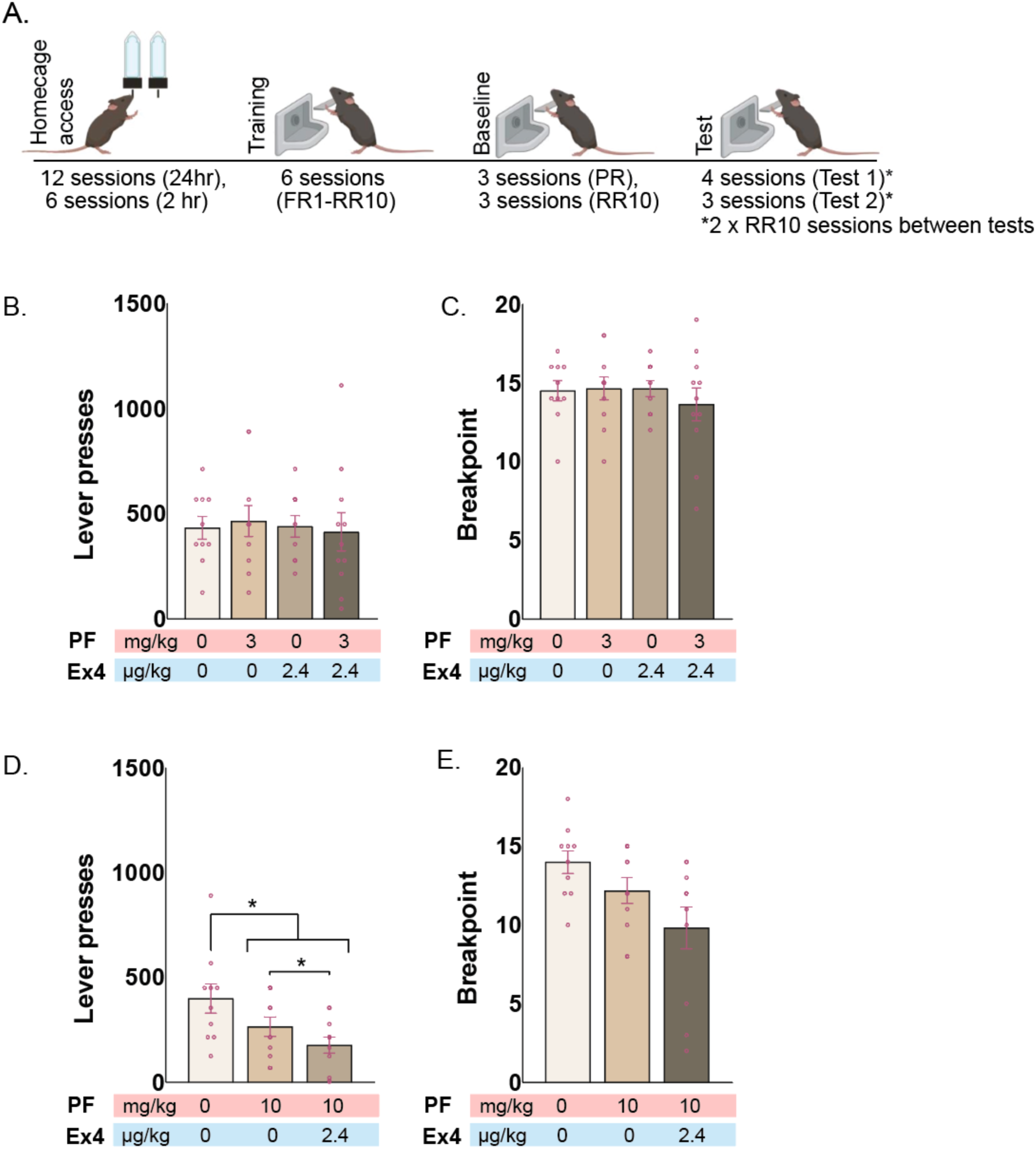
Effects of PF-05231023, Exendin-4, and combination treatment on progressive ratio test of alcohol seeking in male mice (N=12). **(A)** Schematic of alcohol access, training and progressive ratio test. **(B)** Effects of 3mg/kg PF-05231023, 2.4ug/kg Exendin-4, and combination on breakpoint and **(C)** total lever pressing. **(D)** Effects of 10mg/kg PF-05231023 alone and in combination with 2.4ug/kg Exendin-4 on breakpoint and **(E)** total lever pressing. Data are means±SEM. *p<0.05.

During brief 2-hr homecage access, mice consumed 0.92±0.10g/kg (mean±SEM) alcohol solution on average. However, under postprandial conditions (described in supplementary methods), this increased to 2.68±0.51g/kg / 2hr (F(1,9)=18.45, p=.002), confirming the efficacy of this state in enhancing alcohol consumption.

During training, lever-pressing for alcohol increased (F(1,9)=105.041, p<.001). On progressive ratio tests with 10mg/kg PF-05231023, we found reduced motivation for alcohol, indicated by lower breakpoints (Vehicle vs. Treatment: F(1,9)=8.840, p=.016) and fewer lever presses overall (Vehicle vs. Treatment: F(1,9)=7.017, p=.027). Combined PF-05231023/Exendin-4 did not significantly differ from PF-05231023 alone on breakpoints (F(1,9)=3.516, p=.090) but further reduced overall lever pressing relative to PF-05231023 alone (Figs.3D-E).

Notably, PF-05231023’s effects were dose-related. No differences in breakpoint or lever pressing were observed at 3mg/kg for PF-05231023, or in combination with Exendin-4 (p>.60, Figs.3B-C). Exendin-4 was also ineffective when tested alone (F<1). Together these findings suggest enhanced efficacy of PF-05231023 when combined with a subthreshold dose of GLP-1 agonist.

PF-05231023 also dose-dependently reduced 1-hour food intake prior to test (10mg/kg: Vehicle vs. Treatment: F(1,9)=10.250, p=.011; PF-05231023 alone vs combined: p=.526; 3mg/kg: p>.31). Importantly, food reduction at 10mg/kg was unrelated to task performance (Breakpoint: r=−0.478, p=.163; Lever presses: r=−0.251, p=.484), indicating dissociable effects of PF-05231023 on alcohol motivation and food consumption. Finally, no changes were observed in lever-pressing microstructure, including inter-press intervals or post-reward pauses, across doses or treatments (p>.143 for 10mg/kg; p>.089 for 3mg/kg).

### Experiment 4: Cellular imaging of Acb and microstructure of alcohol consumption

We examined how 10mg/kg PF-05231023 affects alcohol consumption by imaging calcium dynamics in Acb neurons during voluntary alcohol intake (20 min sessions; Fig.4A). We assessed lick patterns and corresponding activity of Acb neurons, given that Acb is a critical target through which KLB+ BLA-projecting cells modulate alcohol consumption (24).

**Figure 4.**
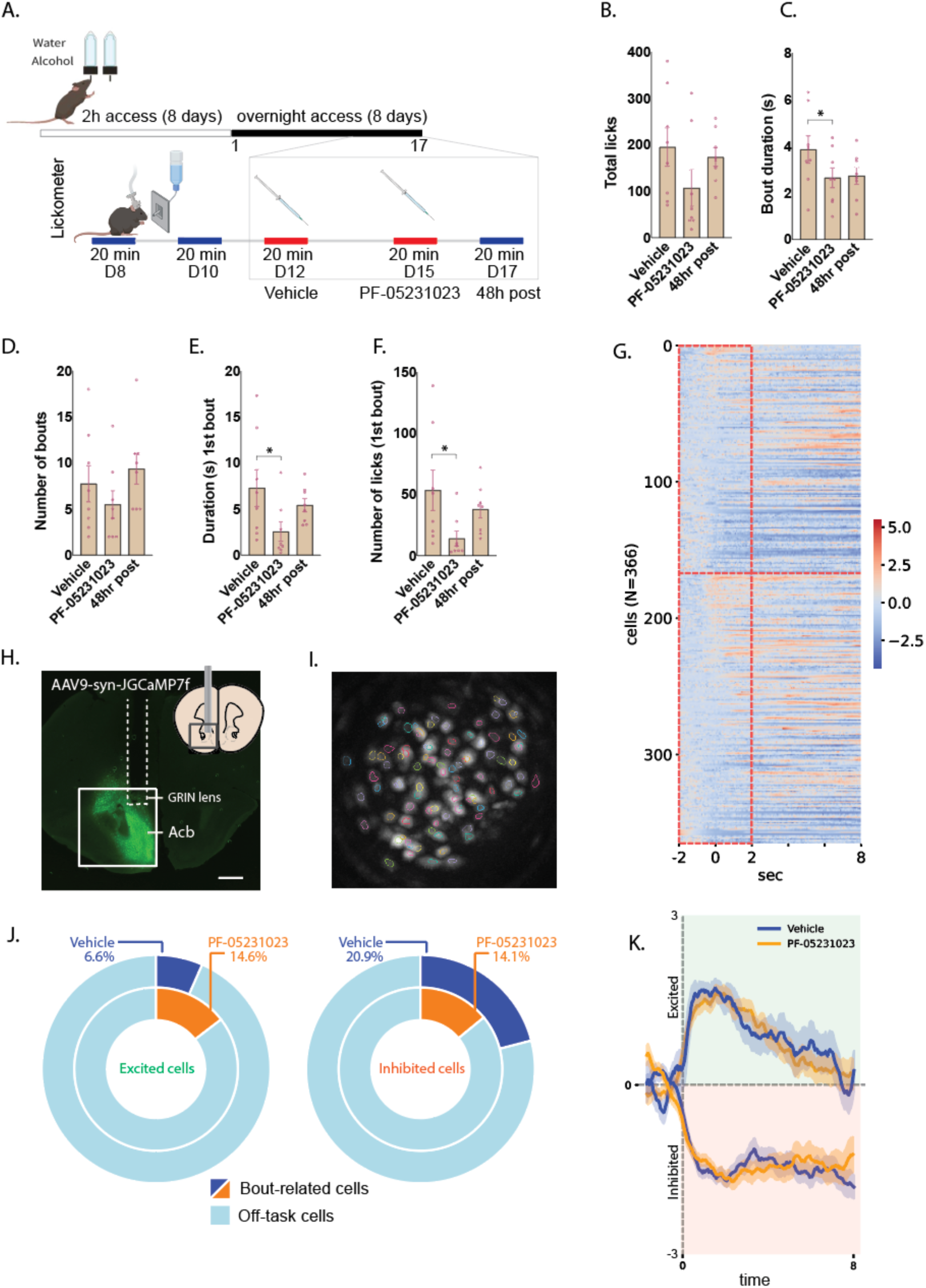
PF-05231023 disrupts lick microstructure during alcohol consumption and modulates consumption-related activity in Acb neurons in male mice (N=8). **(A)** Schematic of training and test procedure. **(B)** Effects of 10mg/kg PF-05231023 on total number of licks, **(C)** bout duration and **(D)** number of bouts across the session; **(E)** Duration and **(F)** Number of licks on the 1^st^ bout. **(G)** Heat map of mean z-scored GCaMP7f peri-bout activity for all neurons during Vehicle (cells 1-199) and PF-05231023 (cells 200-366). **(H)** Example virus expression and GRIN lens position in Acb. **(I)** Example maximum ΔF/F image and identified neurons. **(J)** Fraction and **(K)** Z-score GCaMP7f activity of excited and inhibited neurons. GCaMP7f data are normalised to baseline. Data are means±SEM. *p<0.05.

Mean total licks (±SEM) were 195.1±41.3 (Vehicle), 106.8±39.3 (PF-05231023), and 173.6±20.1 (48-hour post-test, no treatment). While PF-05231023 did not significantly reduce total licks (F(1,7)<1), it disrupted the microstructure of drinking, reducing lick bout duration (Vehicle vs. PF-05231023: F(1,7)=8.17, p=0.024; PF-05231023 vs. 48-hour post: p=0.083) without affecting bout frequency (p>0.05; Figs.4B-D). Effects were more pronounced for the first bout: PF-05231023 reduced both lick count and bout duration (Lick count Vehicle vs. PF-05231023: F(1,7)=7.156, p=.032; Lick count PF-05231023 vs. 48-hr post: F(1,7)=22.466, p=.002; Bout duration Vehicle vs. PF-05231023: F(1,7)=6.648, p=.037; Bout duration PF-05231023 vs. 48-hr post: F(1,7)=10.974, p=.013). PF-05231023 did not alter movement frequency (F(1,7)<1) or subsequent homecage alcohol consumption (F(1,6)<1). The selective effect of PF-05231023 on bout size is consistent with PF-05231023 targeting pre-ingestive evaluative processes (e.g. palatability) (31).

Correspondingly, many Acb neurons exhibited time-locked inhibition at lick bout onset. Under Vehicle, 25.1% (42/167 cells) showed inhibited GCaMP7f transients, while 20.6% (41/199 cells) did so with PF-05231023. Conversely, fewer neurons were excited (Vehicle: 12.0%, 20/167 cells; PF-05231023: 15.1%, 30/199 cells). While there were more inhibited than excited cells under Vehicle (two-proportion Z test, p=0.002), PF-05231023 abolished this imbalance (p=0.15). Analyses of the first lick bout revealed similar trends, with more cells exhibiting inhibition under Vehicle (21%, 35 /167 cells inhibited; 7%, 11/167 cells excited) (p<0.001) but not under PF-05231023 (14%, 28/199 cells inhibited; 15%, 29/199 cells excited; p=0.77, Figs.4J-K). Critically, PF-05231023 did not alter the temporal profiles of bout-related transients (Fig.4K) suggesting that a primary effect of PF-05231023 is to diminish the relative contribution of inhibited cells.

We next assessed cells classified on the basis of their activity around the initiation and termination of a lick bout (ie, bout vs. pause cells). To disentangle bout onset and offset signals, we used an encoding model that removed the linear component of closely-timed events from the target event (32). We found similar proportions of significantly modulated cells between Vehicle (15.0%, 25/167) and PF-05231023 (14.1%, 28/199; Fig.5A). When bout-and pause-encoding cells were separately examined, the percentage of bout cells was also similar across treatments (expressed as a proportion of significantly modulated cells, Vehicle: 84%; PF-05231023: 75%, p>.05, Fig.5B). However, mean kernel GCaMP7f activity of these bout cells was significantly higher following PF-05231023 than Vehicle (F(1,40)=5.251, p=.027; Fig.5C). PF-05231023 also increased the proportion of pause cells (Vehicle: 20%; PF-05231023: 46.4%, p=0.04) without affecting their kernel activity (F<1; Figs.5B-C). Finally, we note that bout-and pause-cell overlap was minimal under Vehicle (4%) but increased with PF-05231023 (21.4%, p=0.06).

**Figure 5.**
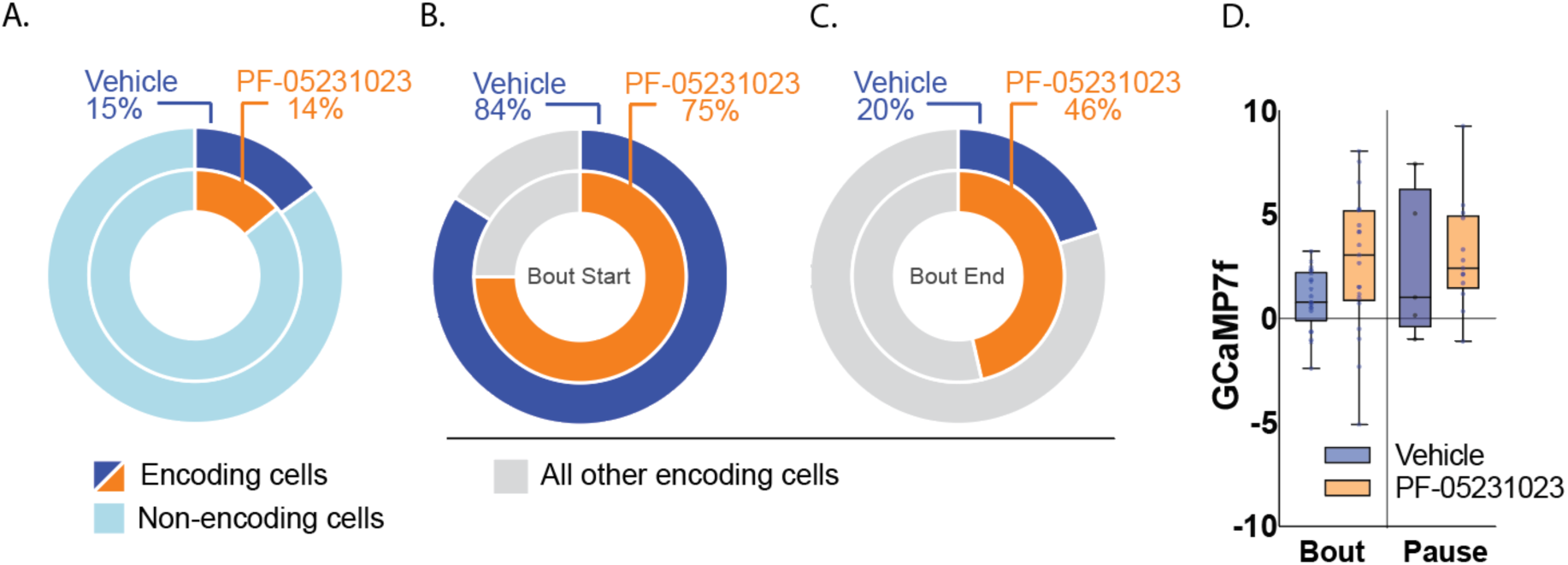
PF-05231023 differentially recruits cells around alcohol consumption in male mice (N=8). **(A)** Fraction of neurons significantly modulated around lick bouts. **(B)** Fraction of bout-modulated cells recruited during bout onset (bout) and **(C)** bout offset (pause). **(D)** Mean GCaMP7f kernel activity over a 2s window following bout onset and offset. *p<0.05.

## Discussion

FGF21 has been shown to regulate alcohol consumption across species (23–27). Here, we demonstrate for the first time its broader influence on alcohol-motivated behaviour and identify potential mechanisms for this regulation, including reducing alcohol palatability and modulating consumption-related Acb neuron activity.

### PF-05231023 and Alcohol Intake

Our findings confirm that PF-05231023 reduces alcohol intake, consistent with prior studies using the same dose in a similar intermittent access model [21]. However, unlike previous reports, we did not observe persistent treatment effects, though we note differences in dosing schedules; we used a single 3mg/kg dose where others have used bi-weekly treatments of the same dose [21]. Notably, we found PF-05231023 reduced alcohol intake only in male mice, contrasting earlier findings of indiscriminate effects of PF-05231023 on alcohol intake across sexes [21].

The cause of this sex difference remains unclear but may relate to reported sex-specific metabolic effects of exogenous FGF21 (14, 33–36), possibly linked to differences in FGF21 receptor expression or liver status following dietary changes (14, 35). In our study, mice had stable and moderate levels of drinking before treatment. Whether similar sex distinctions occur in mice with chronic access to alcohol is not well-characterised but is an important consideration given the hedonic (37) and neurobiological changes associated with prolonged alcohol exposure and the higher alcohol consumption in female mice relative to males.

### FGF21 and Preparatory Appetitive Behaviours

To date, studies have focused on FGF21’s role on consumption but alcohol-associated cues have substantial effects on craving and alcohol-motivated actions. For example, in humans, alcohol cues increase subjective craving, impair inhibitory control, bias attentional processes and increase consumption (38–41). Similarly in rodents, alcohol cues can invigorate instrumental alcohol seeking and drive relapse like behaviour (42–44). The effects of FGF21 on reward-predictive cues have not been previously shown. Here, we show that PF-05231023 had no effect on approach maintained by sucrose cues. This is despite the role for FGF21 in regulating sweet preference (23), and suggests a dissociation in the actions of FGF21 under opportunities of voluntary *ad lib* consumption and cue-controlled approach. Critically, PF-05231023 impaired alcohol approach without reducing the number of trials that elicited approach. This suggests that under these conditions, the primary effect of PF-05231023 may involve reducing the behavioural vigor associated with alcohol anticipation. However, because elevations in approach for alcohol were well-aligned to its availability, further studies are needed to discern the effects of exogenous FGF21 on the incentive and signalling properties of alcohol-related cues.

While the effects of PF-05231023 on alcohol approach are closely tied to the moment of consumption, PF-05231023 also reduced the willingness to work for alcohol on a progressively increasing schedule of reinforcement, a finding that clearly demonstrates the effects of PF-05231023 in a manner temporally dissociated from consumption. Moreover, PF-05231023 disrupted the microstructure of alcohol consumption, specifically reducing the size but not number of lick bouts -a dissociation associated with a reduction in pre-ingestive evaluative processes and reward palatability (31). As such, a change in hedonic responses to alcohol may be one mechanism through which PF-05231023 reduces alcohol intake.

Notably, there was a complex dose-response relationship across our studies. We found that the highest dose was effective on cue-approach, lever-pressing, and lickometer tests, yet ineffective for *ad lib* 24hr homecage consumption, which required a lower dose. This may be due to possible differences in endogenous FGF21 levels at test associated with differences in actual levels of intake between tasks, given that alcohol strongly stimulates hepatic FGF21 release (20, 45, 46); or associated with the food-restricted status of mice undergoing behavioural tests, given that food restriction can elevate FGF21 hepatic expression and plasma levels (47) and that diet induced-elevations in FGF21 can lead to an FGF21 resistant-like state (48). How dose-response functions shift under varying testing conditions warrants further investigation but together, our findings highlight the importance of testing multiple doses when working with FGF21 across different tasks.

### Combined Targeting of FGF21 and GLP-1 reduces alcohol motivation

The broad effects of PF-05231023 across multiple forms of alcohol-motivated behaviour parallel those of gut-brain signalling hormones, like GLP-1, which can reduce self-administration (6, 49, 50), conditioned place preference (6), and relapse (50–52). Critically, our findings suggest positive interactions between GLP-1 and FGF21. Treatment with a sub-threshold dose of the GLP-1 agonist, Exendin-4, augmented the effects of PF-05231023 on reducing total lever presses on a progressive ratio test–a sensitive indicator of motivation over the canonical breakpoint when using an exponential response requirement (53). We note that in contrast to previous studies, Exendin-4, when tested alone, had no effect on instrumental performance (6, 50) which may be due to fasting-dependent changes in GLP-1 receptor localisation or signaling (54).

Together these findings concur with evidence of complimentary and inter-dependent mechanisms of action between FGF21 and GLP-1 (55–58), and recent developments of GLP-1, FGF21 dual agonist compounds (16, 17) in the context of weight loss and insulin sensitivity. Our findings suggest that combination peptide agonist approaches may similarly be of benefit in the treatment of AUD.

### FGF21 Reduces Consumption-Related Pauses in Acb Cells

A wealth of literature implicates overlapping neurobiological substrates mediating FGF21 effects on alcohol consumption and those mediating appetitive behaviour. For example, Pavlovian conditioned approach depends, in part, on glutamatergic projections from basolateral amygdala (BLA) to the Acb (28, 59–62). Optogenetic excitation of this BLA➔Acb pathway can arrest both alcohol consumption and approach to an alcohol cue (28). Crucially, PF-05231023 increases excitability of KLB-expressing BLA➔Acb neurons, and FGF21 suppresses alcohol, but not sucrose, consumption via these BLA KLB+ neurons (24). Our findings further highlights this overlapping role for the Acb. Here we report that distinct subsets of Acb cells are time-locked to lick bouts; that for a majority of these cells, activity is inhibited and that the recruitment of these inhibited cells is undermined by treatment with PF-05231023. Moreover, under PF-05231023, more cells were modulated by the onset of pauses in drinking, consistent with the reduction in bout size on test. Finally, while PF-05231023 did not significantly alter the number of cells recruited during the onset of a lick bout, it increased the activity of these cells. These findings fit within an established role for Acb in the control over consumption and learned appetitive behaviour (29, 30, 63–65). Moreover, they suggest that a retuning of specific subsets of Acb cells associated with initiating and terminating consumption may be a key mechanism through which PF-05231023 exerts control over alcohol intake; a theory that invites further scrutiny.

### Conclusions

PF-05231023 reduces alcohol consumption, approach, and motivation, likely through reductions in hedonic responses to alcohol and via modulation of Acb neuron activity to control intake. These findings highlight FGF21’s potential as a therapeutic target for AUD.

## Supporting information

Supplemental Methods and Results

## Acknowledgements

Supported by a Synergy Grant from the National Health and Medical Research Council of Australia (GNT 2009851) and BBRF NARSAD Young Investigator Award

## Data Availability Statement

Data will be shared and made available upon reasonable request. Code and further information related to the event-encoding model will be shared upon reasonable request and made publicly available on GitHub as of the date of publication.

## Author Contributions

BJC, AJL, GPM, PSH, and EZM conceived and designed the work and wrote the original draft. BJC, CVO H-B, WAH, and AWL contributed to data collection. CVO H-B and EAC contributed to miniscope data collection. BJC, CVO H-B, GPM and EZM analysed the data. CVO H-B, GPM, and EZM analysed the miniscope data. GPM, AJL, PSH and EZM revised the manuscript.

## References

1. Global status report on alcohol and health and treatment of substance use disorders. Geneva, 2024.

2. Kranzler HR, Soyka M. Diagnosis and Pharmacotherapy of Alcohol Use Disorder: A Review. Jama. 2018;320(8):815–24.

3. McPheeters M, O’Connor EA, Riley S, Kennedy SM, Voisin C, Kuznacic K, et al. Pharmacotherapy for Alcohol Use Disorder: A Systematic Review and Meta-Analysis. Jama. 2023;330(17):1653–65.

4. Leggio L, Hendershot CS, Farokhnia M, Fink-Jensen A, Klausen MK, Schacht JP, et al. GLP-1 receptor agonists are promising but unproven treatments for alcohol and substance use disorders. Nat Med. 2023;29(12):2993–5.

5. Klausen MK, Jensen ME, Møller M, Le Dous N, Jensen A-MØ, Zeeman VA, et al. Exenatide once weekly for alcohol use disorder investigated in a randomized, placebo-controlled clinical trial. JCI Insight. 2022;7(19).

6. Egecioglu E, Steensland P, Fredriksson I, Feltmann K, Engel JA, Jerlhag E. The glucagon-like peptide 1 analogue Exendin-4 attenuates alcohol mediated behaviors in rodents. Psychoneuroendocrinology. 2013;38(8):1259–70.

7. Farokhnia M, Grodin EN, Lee MR, Oot EN, Blackburn AN, Stangl BL, et al. Exogenous ghrelin administration increases alcohol self-administration and modulates brain functional activity in heavy-drinking alcohol-dependent individuals. Mol Psychiatry. 2018;23(10):2029–38.

8. Lei K, Wegner SA, Yu J-H, Hopf FW. Orexin-1 receptor blockade suppresses compulsive-like alcohol drinking in mice. Neuropharmacology. 2016;110:431–7.

9. Flores-Ramirez FJ, Illenberger JM, Pascasio GE, Matzeu A, Mason BJ, Martin-Fardon R. Alternative use of suvorexant (Belsomra®) for the prevention of alcohol drinking and seeking in rats with a history of alcohol dependence. Frontiers in Behavioral Neuroscience. 2022;16.

10. Campbell EJ, Bonomo Y, Collins L, Norman A, O’Neill H, Streitberg A, et al. The dual orexin receptor antagonist suvorexant in alcohol use disorder and comorbid insomnia: A case report. Clin Case Rep. 2024;12(5):e8740.

11. Bach P, Bumb JM, Schuster R, Vollstädt-Klein S, Reinhard I, Rietschel M, et al. Effects of leptin and ghrelin on neural cue-reactivity in alcohol addiction: Two streams merge to one river? Psychoneuroendocrinology. 2019;100:1–9.

12. Hillemacher T, Kraus T, Rauh J, Weiß J, Schanze A, Frieling H, et al. Role of Appetite-Regulating Peptides in Alcohol Craving: An Analysis in Respect to Subtypes and Different Consumption Patterns in Alcoholism. Alcoholism: Clinical and Experimental Research. 2007;31(6):950–4.

13. Solon-Biet SM, Clark X, Bell-Anderson K, Rusu PM, Perks R, Freire T, et al. Toward reconciling the roles of FGF21 in protein appetite, sweet preference, and energy expenditure. Cell Reports. 2023;42(12):113536.

14. Zhou B, Claflin KE, Flippo KH, Sullivan AI, Asghari A, Tadinada SM, et al. Central FGF21 production regulates memory but not peripheral metabolism. Cell Reports. 2022;40(8).

15. Hill CM, Laeger T, Dehner M, Albarado DC, Clarke B, Wanders D, et al. FGF21 Signals Protein Status to the Brain and Adaptively Regulates Food Choice and Metabolism. Cell Rep. 2019;27(10):2934–47.e3.

16. Gilroy CA, Capozzi ME, Varanko AK, Tong J, D’Alessio DA, Campbell JE, et al. Sustained release of a GLP-1 and FGF21 dual agonist from an injectable depot protects mice from obesity and hyperglycemia. Sci Adv. 2020;6(35):eaaz9890.

17. Pan Q, Lin S, Li Y, Liu L, Li X, Gao X, et al. A novel GLP-1 and FGF21 dual agonist has therapeutic potential for diabetes and non-alcoholic steatohepatitis. EBioMedicine. 2021;63:103202.

18. Farokhnia M, Wang T, Jourdan T, Godlewski G, Farinelli LA, Kunos G, et al. A human laboratory study on the link between alcohol administration and circulating fibroblast growth factor 21 (FGF21) in individuals with alcohol use disorder. Drug Alcohol Depend. 2023;245:109809.

19. Chen Z, Yang L, Liu Y, Huang P, Song H, Zheng P. The potential function and clinical application of FGF21 in metabolic diseases. Front Pharmacol. 2022;13:1089214.

20. Lanng AR, Gasbjerg LS, Bergmann NC, Bergmann S, Helsted MM, Gillum MP, et al. Gluco-metabolic effects of oral and intravenous alcohol administration in men. Endocr Connect. 2019;8(10):1372–82.

21. Shi SY, Lu Y-W, Richardson J, Min X, Weiszmann J, Richards WG, et al. A systematic dissection of sequence elements determining β-Klotho and FGF interaction and signaling. Scientific Reports. 2018;8(1):11045.

22. Kurosu H, Choi M, Ogawa Y, Dickson AS, Goetz R, Eliseenkova AV, et al. Tissue-specific expression of betaKlotho and fibroblast growth factor (FGF) receptor isoforms determines metabolic activity of FGF19 and FGF21. J Biol Chem. 2007;282(37):26687–95.

23. Talukdar S, Owen BM, Song P, Hernandez G, Zhang Y, Zhou Y, et al. FGF21 Regulates Sweet and Alcohol Preference. Cell Metab. 2016;23(2):344–9.

24. Flippo KH, Trammell SAJ, Gillum MP, Aklan I, Perez MB, Yavuz Y, et al. FGF21 suppresses alcohol consumption through an amygdalo-striatal circuit. Cell Metab. 2022;34(2):317–28.e6.

25. Clarke TK, Adams MJ, Davies G, Howard DM, Hall LS, Padmanabhan S, et al. Genome-wide association study of alcohol consumption and genetic overlap with other health-related traits in UK Biobank (N=112 117). Mol Psychiatry. 2017;22(10):1376–84.

26. Ho MF, Zhang C, Moon I, Wei L, Coombes B, Biernacka J, et al. Genome-wide association study for circulating FGF21 in patients with alcohol use disorder: Molecular links between the SNHG16 locus and catecholamine metabolism. Mol Metab. 2022;63:101534.

27. Schumann G, Liu C, O’Reilly P, Gao H, Song P, Xu B, et al. KLB is associated with alcohol drinking, and its gene product β-Klotho is necessary for FGF21 regulation of alcohol preference. Proc Natl Acad Sci U S A. 2016;113(50):14372–7.

28. Millan EZ, Kim HA, Janak PH. Optogenetic activation of amygdala projections to nucleus accumbens can arrest conditioned and unconditioned alcohol consummatory behavior. Neuroscience. 2017;360:106–17.

29. Krause M, German PW, Taha SA, Fields HL. A pause in nucleus accumbens neuron firing is required to initiate and maintain feeding. J Neurosci. 2010;30(13):4746–56.

30. O’Connor EC, Kremer Y, Lefort S, Harada M, Pascoli V, Rohner C, et al. Accumbal D1R Neurons Projecting to Lateral Hypothalamus Authorize Feeding. Neuron. 2015;88(3):553–64.

31. Johnson AW. Characterizing ingestive behavior through licking microstructure: Underlying neurobiology and its use in the study of obesity in animal models. Int J Dev Neurosci. 2018;64:38–47.

32. Parker NF, Baidya, A., Cox, J., Haetzel, L. M., Zhukovskaya, A., Murugan, M., Engelhard, B., Goldman, M. S., & Witten, I. B.. Choice-selective sequences dominate in cortical relative to thalamic inputs to NAc to support reinforcement learning. Cell reports,. 2022;39(7).

33. Chaffin AT, Larson KR, Huang KP, Wu CT, Godoroja N, Fang Y, et al. FGF21 controls hepatic lipid metabolism via sex-dependent interorgan crosstalk. JCI Insight. 2022;7(19).

34. Makarova E, Kazantseva A, Dubinina A, Jakovleva T, Balybina N, Baranov K, et al. The Same Metabolic Response to FGF21 Administration in Male and Female Obese Mice Is Accompanied by Sex-Specific Changes in Adipose Tissue Gene Expression. Int J Mol Sci. 2021;22(19).

35. Makarova EN, Yakovleva TV, Balyibina N, Baranov KO, Denisova EI, Dubinina A, et al. Pharmacological effects of fibroblast growth factor 21 are sex-specific in mice with the lethal yellow (Ay) mutation. Vavilov Journal of Genetics and Breeding. 2020;24:200–8.

36. Soto Sauza KA, Ryan KK. FGF21 mediating the Sex-dependent Response to Dietary Macronutrients. J Clin Endocrinol Metab. 2024;109(9):e1689–e96.

37. Bachmanov AA, Kiefer SW, Molina JC, Tordoff MG, Duffy VB, Bartoshuk LM, et al. Chemosensory factors influencing alcohol perception, preferences, and consumption. Alcohol Clin Exp Res. 2003;27(2):220–31.

38. Chen K, Garbusow M, Sebold M, Kuitunen-Paul S, Smolka MN, Huys QJM, et al. Alcohol Approach Bias Is Associated With Both Behavioral and Neural Pavlovian-to-Instrumental Transfer Effects in Alcohol-Dependent Patients. Biological Psychiatry Global Open Science. 2023;3(3):443–50.

39. Courtney KE, Ray LA. Subjective responses to alcohol in the lab predict neural responses to alcohol cues. J Stud Alcohol Drugs. 2014;75(1):124–35.

40. Field M, Jones A. Elevated alcohol consumption following alcohol cue exposure is partially mediated by reduced inhibitory control and increased craving. Psychopharmacology (Berl). 2017;234(19):2979–88.

41. Jasinska AJ, Stein EA, Kaiser J, Naumer MJ, Yalachkov Y. Factors modulating neural reactivity to drug cues in addiction: a survey of human neuroimaging studies. Neurosci Biobehav Rev. 2014;38:1–16.

42. Alarcón DE, Delamater AR. Outcome-specific Pavlovian-to-instrumental transfer (PIT) with alcohol cues and its extinction. Alcohol. 2019;76:131–46.

43. Corbit LH, Janak PH. Habitual Alcohol Seeking: Neural Bases and Possible Relations to Alcohol Use Disorders. Alcohol Clin Exp Res. 2016;40(7):1380–9.

44. Katner SN, Magalong JG, Weiss F. Reinstatement of alcohol-seeking behavior by drug-associated discriminative stimuli after prolonged extinction in the rat. Neuropsychopharmacology. 1999;20(5):471–9.

45. Desai BN, Singhal G, Watanabe M, Stevanovic D, Lundasen T, Fisher FM, et al. Fibroblast growth factor 21 (FGF21) is robustly induced by ethanol and has a protective role in ethanol associated liver injury. Mol Metab. 2017;6(11):1395–406.

46. Søberg S, Andersen ES, Dalsgaard NB, Jarlhelt I, Hansen NL, Hoffmann N, et al. FGF21, a liver hormone that inhibits alcohol intake in mice, increases in human circulation after acute alcohol ingestion and sustained binge drinking at Oktoberfest. Mol Metab. 2018;11:96–103.

47. Badman MK, Pissios P, Kennedy AR, Koukos G, Flier JS, Maratos-Flier E. Hepatic fibroblast growth factor 21 is regulated by PPARalpha and is a key mediator of hepatic lipid metabolism in ketotic states. Cell Metab. 2007;5(6):426–37.

48. Fisher FM, Chui PC, Antonellis PJ, Bina HA, Kharitonenkov A, Flier JS, et al. Obesity is a fibroblast growth factor 21 (FGF21)-resistant state. Diabetes. 2010;59(11):2781–9.

49. Chuong V, Farokhnia M, Khom S, Pince CL, Elvig SK, Vlkolinsky R, et al. The glucagon-like peptide-1 (GLP-1) analogue semaglutide reduces alcohol drinking and modulates central GABA neurotransmission. JCI Insight. 2023;8(12).

50. Díaz-Megido C, Thomsen M. Sex-dependent divergence in the effects of GLP-1 agonist exendin-4 on alcohol reinforcement and reinstatement in C57BL/6J mice. Psychopharmacology. 2023;240(6):1287–98.

51. Aranäs C, Edvardsson CE, Shevchouk OT, Zhang Q, Witley S, Blid Sköldheden S, et al. Semaglutide reduces alcohol intake and relapse-like drinking in male and female rats. EBioMedicine. 2023;93:104642.

52. Thomsen M, Dencker D, Wörtwein G, Weikop P, Egecioglu E, Jerlhag E, et al. The glucagon-like peptide 1 receptor agonist Exendin-4 decreases relapse-like drinking in socially housed mice. Pharmacol Biochem Behav. 2017;160:14–20.

53. Tsibulsky VL, Norman AB. Methodological and analytical issues of progressive ratio schedules: Definition and scaling of breakpoint. J Neurosci Methods. 2021;356:109146.

54. Ronveaux CC, de Lartigue G, Raybould HE. Ability of GLP-1 to decrease food intake is dependent on nutritional status. Physiol Behav. 2014;135:222–9.

55. Le TDV, Fathi P, Watters AB, Ellis BJ, Besing GK, Bozadjieva-Kramer N, et al. Fibroblast growth factor-21 is required for weight loss induced by the glucagon-like peptide-1 receptor agonist liraglutide in male mice fed high carbohydrate diets. Mol Metab. 2023;72:101718.

56. Liu J, Yang K, Yang J, Xiao W, Le Y, Yu F, et al. Liver-derived fibroblast growth factor 21 mediates effects of glucagon-like peptide-1 in attenuating hepatic glucose output. EBioMedicine. 2019;41:73–84.

57. Liu D, Pang J, Shao W, Gu J, Zeng Y, He HH, et al. Hepatic Fibroblast Growth Factor 21 Is Involved in Mediating Functions of Liraglutide in Mice With Dietary Challenge. Hepatology. 2021;74(4):2154–69.

58. Yang M, Zhang L, Wang C, Liu H, Boden G, Yang G, et al. Liraglutide Increases FGF-21 Activity and Insulin Sensitivity in High Fat Diet and Adiponectin Knockdown Induced Insulin Resistance. PLOS ONE. 2012;7(11):e48392.

59. Di Ciano P, Cardinal RN, Cowell RA, Little SJ, Everitt BJ. Differential involvement of NMDA, AMPA/kainate, and dopamine receptors in the nucleus accumbens core in the acquisition and performance of pavlovian approach behavior. J Neurosci. 2001;21(23):9471–7.

60. Di Ciano P, Everitt BJ. Direct interactions between the basolateral amygdala and nucleus accumbens core underlie cocaine-seeking behavior by rats. J Neurosci. 2004;24(32):7167–73.

61. Valyear MD, Brown A, Deyab G, Villaruel FR, Lahlou S, Caporicci-Dinucci N, et al. Augmenting glutamatergic, but not dopaminergic, activity in the nucleus accumbens shell disrupts responding to a discrete alcohol cue in an alcohol context. European Journal of Neuroscience. 2024;59(7):1500–18.

62. Puaud M, Higuera-Matas A, Brunault P, Everitt BJ, Belin D. The Basolateral Amygdala to Nucleus Accumbens Core Circuit Mediates the Conditioned Reinforcing Effects of Cocaine-Paired Cues on Cocaine Seeking. Biological Psychiatry. 2021;89(4):356–65.

63. Maldonado-Irizarry CS, Swanson CJ, Kelley AE. Glutamate receptors in the nucleus accumbens shell control feeding behavior via the lateral hypothalamus. J Neurosci. 1995;15(10):6779–88.

64. Taha SA, Fields HL. Encoding of palatability and appetitive behaviors by distinct neuronal populations in the nucleus accumbens. J Neurosci. 2005;25(5):1193–202.

65. Prado L, Luis-Islas J, Sandoval OI, Puron L, Gil MM, Luna A, et al. Activation of Glutamatergic Fibers in the Anterior NAc Shell Modulates Reward Activity in the aNAcSh, the Lateral Hypothalamus, and Medial Prefrontal Cortex and Transiently Stops Feeding. J Neurosci. 2016;36(50):12511–29.

