## Supplemental Methods and Results for "FGF21 Analogue PF-05231023 on Alcohol Consumption and Neuronal Activity in the Nucleus Accumbens"

### Supplementary Material

#### Materials and Methods

*Surgery*

Mice were deeply anaesthetised with 5% v/v isoflurane in oxygen-enriched air after subcutaneous injection of 5 mg/kg carprofen (Rimadyl, Zoetis). They were secured into a stereotaxic frame (Model 1900, Kopf Instruments) and maintained on 1%-2.5% v/v isoflurane for the duration of surgery. Before scalp incision, a local anaesthetic was delivered to the incision site (0.1 ml Marcaine (0.5%), s.c.). Ophthalmic gel (Viscotears, Alcon) was applied to avoid eye drying.

To track dynamic changes in the neural activity of individual Acb cells, a calcium sensor packaged in an adeno-associated virus (AAV9-syn-JGCaMP7f-WPRE, Addgene, ~2.6 x $10^{13}$ (viral particles), 0.5ul injected at 1:3 dilution in PBS) was delivered into the right or left Acb (+1.42 A-P, +/-0.5 M-L, -4.35 and –4.25 D-V) using a 33-gauge syringe (Hamilton) via a microinjection pump (World Precision Instruments). A retrograde adeno-associated virus encoding the fluorescent label, mCherry (AAVrg-hsyn-MCherry, Addgene, ~2.6 x $10^{13}$ (viral particles), 0.2ul) was delivered into the ipsilateral lateral hypothalamus (LH; -1.4 A-P, +/-1.25 M-L, -5.2 D-V) to label LH-projecting Acb cells. However, there was no signal detected under the miniscope and histological examination confirmed all LH-injections were misplaced and/or poorly-expressed. Therefore, we did not include tracing data in our analyses. Mice were also implanted with a 0.5mm gradient refractive index (GRIN) lens (GLP-0561, Inscopix) at the same Acb A-P and M-L coordinate, dorsally-position (-4.25 D-V), which was secured to the skull with dental adhesive and miniaturised stainless-steel screws. Mice received the antibiotic Duplocillin (0.15 ml/kg, s.c.) immediately after surgery and were closely monitored during recovery. 2-3 weeks post-surgery, mice were anaesthetised with 5% v/v isoflurane in oxygen-enriched air, placed in the stereotaxic frame as described above, and a base plate attached to a miniature one-photon detachable microscope (miniscope, nVue system, Inscopix) was secured above the implanted lens at a distance with a focused field of view with dental adhesive.

All coordinates are relative to bregma using the Paxinos and Franklin's Mouse Brain in Stereotaxic Coordinates [28].

**Behavioural Apparatus**

*Home cage two-bottle choice consumption.* We used open-source 3D printed sippers to detect consummatory behaviours during homecage alcohol consumption [28]. Briefly, the sippers housed an Adafruit Feather M0 Adalogger microcontroller, battery and infrared photo-interrupters detecting interactions with the sipper (frequency and duration). Bottles were 15ml centrifuge tubes fitted with Hydropac valves (Allied Scientific Products, Australia), sealed with rubber O rings.

*Behavioural training.* We used standard Med Associates chambers. with clear Perspex walls and stainless-steel grid floors enclosed in light and sound attenuating cabinets. Each box contained a speaker and central recessed magazine. For alcohol cue studies, the chambers contained a pellet dispenser and dipper arm for delivering 15% v/v alcohol in a 10cc cup. For operant studies, the chambers contained a retractable lever on the left or right side of a magazine. Alcohol was delivered into the magazine via silicone tubing fitted to a 60mL syringe on a syringe pump. Magazine entries were detected by an infrared photobeam. For lickometer studies, mice accessed alcohol solution (15% v/v) via a stainless-steel spout protruding from one side-wall of the chamber. Licks were detected via closed-loop contact between the spout and grid floor. All experimental events were controlled by Med-PC and Med Associates software.

*Cellular-resolution calcium imaging.* Mice were habituated to the imaging procedure prior to training over three days. The miniscope was attached and mice were subsequently placed in a 20-litre polypropylene bucket lined with corn cob bedding for 15 minutes.

To capture in vivo cellular imaging of Acb cells during alcohol consumption, lick-event timestamps were synchronised to the neural recordings via lick-paired TTL signals from Med Associates to the DAQ. LED illumination emitted from the miniscope was triggered at the start of each session, controlled similarly by a Med Associates input to the DAQ. GCaMP7f fluorescence was captured under the blue (455nm) excitation wavelength at 20 frames/second. Recording parameters were held constant across mice using Inscopix software (Data Acquisition Software; spatial downsampling = 2; temporal downsampling = 0; gain = 6.0; LED = 0.03mW $mm^{-2}$).

Acquired movies were then batch-processed through an Inscopix Data Processing pipeline using a custom Application Programming Interface (API) in Python. Specific modules were provided in the Inscopix API package (https://inscopix.github.io/pyisx/reference.html). For each movie, field of view and estimated cell diameters were manually determined prior to processing. Each movie file underwent spatial bandpass filtering to remove noise and out-of-focus cells, presenting as low and high spatial frequency content, respectively; motion correction to hold cell locations constant across frames; and cell identification using the extended constrained nonnegative matrix factorization algorithm (CNMF-E, [29]). Results were visually inspected for cell yield, duplication and over-segmentation. Motion-corrected files were iterated through the CNMF-E process with adjusted threshold parameters (merge threshold, min_corr and min_pnr) as needed to optimise cell detection. Cell duplicates were manually rejected. Fluorescence traces of accepted cells from 8 mice (referred to as 'GCaMP7f') were pooled and used for all subsequent analyses.

**Procedure**

*Home cage two-bottle choice consumption.* Mice had access to two bottles (15% v/v alcohol and tapwater) on an intermittent schedule (3 days/week) [30]. One bottle contained 15% v/v alcohol and the other tap water. Bottle positions were alternated to prevent side preference. Intake was measured by weighing bottles before and after each session.

*Conditioned Approach Task.* Mice were trained to associate auditory cues with alcohol (CS1) or sucrose (CS2). They received a 20-minute magazine training session comprising 15 deliveries of 20 mg sucrose pellets (Bio-Serv; VI60). Then, 16 daily sucrose cue training sessions (30 trials: 15 CS+, 15 CS; pseudorandom order; VI range 100-120s). A 10 second tone (2.5 kHz or 10 kHz, counterbalanced) served as the CS+ that co-terminated with delivery of one 20mg sucrose pellet into the magazine. A 10 second white noise served as the CS- and had no consequence. Following training, mice received 3 weeks of overnight intermittent access to alcohol. They then received 16 daily alcohol cue training sessions that proceeded in the same manner as sucrose cue training with the exceptions that a different tone (2.5 kHz or 10 kHz) served as the CS+ and alcohol served as the reinforcer. At the conclusion of training, mice received 1 day of sucrose cue retraining.

*Progressive ratio*

Mice received 4 weeks of 24-hr homecage intermittent access to alcohol followed by 2 weeks of 2-hour intermittent access (1300 – 1500 hrs). Next, they were trained in operant boxes to self-administer alcohol on an intermittent schedule (3 days / week). They received one 15-minute habituation to the operant boxes followed by two 20-minute sessions of magazine training where 0.04ml alcohol was delivered on a variable schedule (VI-60). They then received alcohol-reinforced lever press training under the following sequence of fixed (FR) and random (RR) reinforcement schedules: FR1, RR2, RR5, and RR10 (minimum of two sessions on RR5 and RR10). Mice were required to earn 15 rewards to progress to the next criterion. The session ended after mice received 20 ethanol deliveries or after 40 minutes lapsed, whichever occurred first. Following training, lever pressing was reinforced under an exponentially increasing schedule of reinforcement (i.e. progressive ratio; PR); the number of presses required to deliver alcohol followed the function: $5e^{(\text{alcohol reinforcement} \times 0.2)} - 5$ (i.e., 1, 2, 4, 6, 9, 12, 15, 20, 25, 32, 40, 50, 62, 77, 95, 118, 145... 603 presses) [1]. The session ended following 5 minutes with no lever press or at 3 hours, whichever occurred first. Mice received 3 baseline sessions under a PR schedule followed by 3 retraining (RR10) sessions. Across days of training and on test, mice received 1hr *ad lib* access to food with no water immediately prior to placement in the testing chamber, exploiting the postprandial state to encourage alcohol seeking and intake on this task [2].

*Cellular recordings during alcohol consumption*

Mice received the following intermittent homecage alcohol access schedule: 2hr access for 8 days then 24hr access for 12 days. Starting on the fourth day of access and on subsequent days of access, mice were allowed 20 minutes to freely consumed alcohol in a lickometer chamber. At the conclusion of test, mice received 16-hour homecage alcohol access (starting 17:30). Recordings of Acb task-related cellular activity were acquired on Days 5-8. They received saline to acclimatise to the injection procedure on Day 5, then Vehicle and PF-05231023 (10mg/kg) on Days 6 and 7.

Prior to this study alcohol-naïve mice received alcohol cue training whereby a 10 second auditory cue served as a conditioning stimulus (CS+, 5kHz tone, VI-40 intertrial interval), which preceded the delivery of small drop of alcohol (25ul, 2.5s infusion), 8 seconds following the onset of the CS+. Mice received 10 CS+ presentations in a 20-minute session over 16 days. Recordings of Acb task-related cellular activity were acquired on Days 17-21. They received saline, vehicle and PF-05231023 (10mg/kg) on Days 18-20, respectively. They were retrained (4 days) then tested and again on Days 25-28 where they received saline, vehicle and PF-05231023 (5mg/kg) on Days 25-27, respectively. However, on the first test block, Vehicle had unexpected suppressive effects on consumption intake, which rendered the results uninterpretable. For this reason, we did not analyse this data further and switched to a more direct test of consumption.

**Data analyses and statistics**

*Homecage sippers.* Long entry durations (>15 seconds) were excluded from analyses based on pilot studies where we determined that these long entry times were likely caused by residual liquid on the sipper triggering the photodetectors after the mouse had exited. Data from three mice were excluded due to sipper malfunction on test. We validated the sipper model by performing a correlation and linear regression using the 'statsmodels' package in Python on the percent preference for the alcohol sipper (frequency, duration) vs percentage preference for alcohol consumption across the four test days. We analysed sipper data on test using similar correlations and linear regressions.

*Cue-response.* Mice discriminated between the rewarded (CS+) and unrewarded (CS-) cue when reinforced with sucrose but not alcohol. As such we report conditioned approach (normalised to a 10 second preCS baseline) only for sucrose cues and separately, an aggregate cue responsivity measure (the sum of responding to CS+ and CS-, normalised to a 10 second preCS baseline) for tests involving alcohol. Two animals (n = 2 female) were excluded from sucrose analyses due to poor discrimination in the conditioned approach at the end of training (CS+ - CS- </=0). Cue-related responses were analysed using a within-subjects factor for event (CS+, CS- for cue discrimination; preCS, CS for cue reactivity) and Drug (VEH, PF-05231023). Data were analysed with sexes pooled because we did not have sufficient numbers to examine males (n=6) and females (n=4) separately.

*Progressive ratio test.* We measured total lever presses and breakpoint as the total number of completed response requirements (or total rewards received). We applied log transformation to epoch data including duration of inter-press interval and post-reinforcement pause. One mouse failed to consume food during the 1hour pre-test phase on the vehicle test day and was excluded from analyses.

*Lick response, head acceleration and calcium imaging.* Lick bouts were defined as 3 successive licks with less than 1s inter-lick interval. We normalised GCaMP7f activity to the median and standardised the scores against the median absolute deviation (MAD) from the median score for GCaMP7f recordings of individual cells across the entire recording session.

Mean peri-event GCaMP7f activity for each cell was inspected across a 4 second window (2 seconds before and after lick bout onset). We refer to cells with average population weighted Z-scored GCaMP7f activity exceeding 1 Z-score as 'excited' and those below –1 Z as 'inhibited'. To select cells that were significantly modulated by bout-related events, we generated an estimate (i.e. kernel) of the bout-related activity (e.g. bout onset) of a given cell while accounting for the linear component of the response of this cell to temporally-proximal events (e.g. bout offset). Normalised GCaMP7f data from each cell was fed through a multilinear encoding model to generate the response kernel for each behavioral event [3]. In this model, the dependent variable was the GCaMP7f activity of a single cell during the entire session. The predictors were times of each consumption bout event (bout onset, bout offset), where a bout was defined as a minimum of 3 licks with no more than 1 second between each lick. They were each convolved with a spline basis set generated in RStudio (splines2 package, degrees of freedom = 25) that spanned 2 seconds before to 2 second after the event. This generated 25 temporally delayed versions of the events to predict neural activity. Using the regression coefficient generated from the model, we then calculated a response kernel for each event.

Head acceleration was detected via the 3-axis (xyz) accelerometer built into the nVue camera and acquired using Inscopix software (Data Acquisition Software). Acceleration data was recorded at 50 Hz temporal resolution and downsampled to 10Hz. Head acceleration was defined as the rectified sum of acceleration in all axes, filtered at 5 Hz with a zero-phase third order Butterworth filter, from the accelerometer position. Due to the unimodal distribution of the movement data, we applied K means clustering to determine a threshold; values above this threshold were considered movement events and values below were considered as rest.

### Supplementary Results

### S1.

**Table 1.** Mean±SEM (g/kg) 24-hour home cage alcohol consumption on the last 3 days of access

| | Overall | Males | Females |
|---|---|---|---|
| Experiment 1 | 9.19 ± 0.56 | 6.86 ± 0.43 | 11.38 ± 0.84 |
| Experiment 2 | 8.95 ± 0.77 | 7.41 ± 0.93 | 10.23 ± 1.11 |
| Experiment 3 | 5.62 ± 0.36 | 5.62 ± 0.36g | NA |
| Experiment 4 | 8.42 ± 1.16 | 8.42 ± 1.16 | NA |

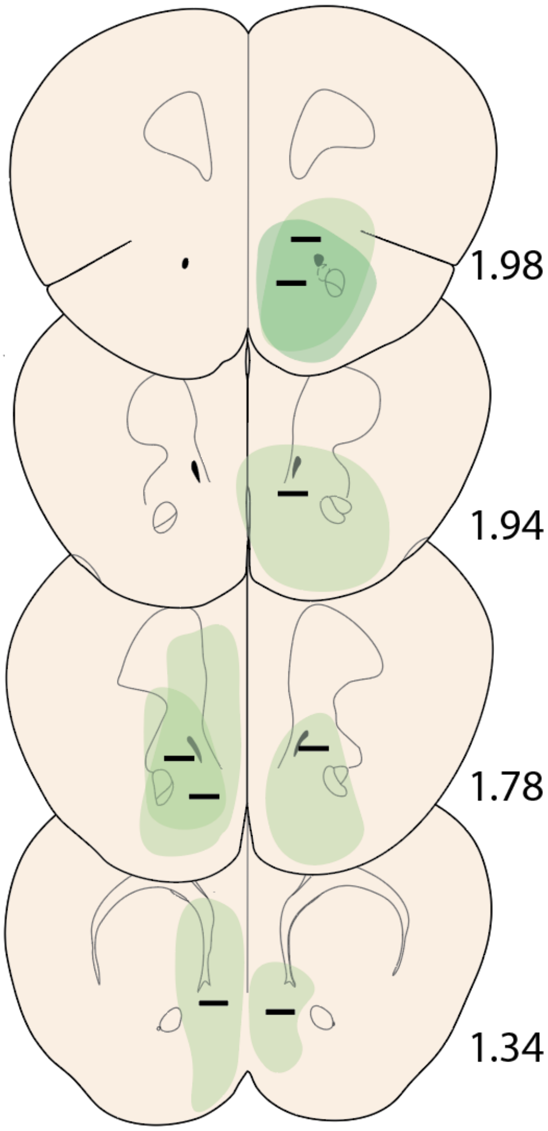


**S2.**

Schematic depicting histological verification of the positioning of the GRIN lens and spread of the Ca2+ sensor, GCaMP7f for mice in Experiment 4 (N=8). Numbers are anterior-posterior (AP) coordinates relative to bregma.

**References**


1. Richardson, N.R. and D.C.S. Roberts, *Progressive ratio schedules in drug self-administration studies in rats: a method to evaluate reinforcing efficacy*. Journal of Neuroscience Methods, 1996. **66**(1): p. 1-11.
2. Middaugh, L.D., et al., *Naltrexone effects on ethanol reward and discrimination in C57BL/6 mice*. Alcohol Clin Exp Res, 1999. **23**(3): p. 456-64.

3. Parker, N.F., Baidya, A., Cox, J., Haetzel, L. M., Zhukovskaya, A., Murugan, M., Engelhard, B., Goldman, M. S., & Witten, I. B. , *Choice-selective sequences dominate in cortical relative to thalamic inputs to NAc to support reinforcement learning*. Cell reports,, 2022. **39**(7).
